# Focusing on what matters: Modulation of the human hippocampus by relational attention

**DOI:** 10.1101/446443

**Authors:** Natalia I. Córdova, Nicholas B. Turk-Browne, Mariam Aly

**Author notes:** Please address correspondence to: Mariam Aly, Department of Psychology, Columbia University, 406 Schermerhorn Hall, 1190 Amsterdam Avenue, New York, NY 10027.

## Abstract

Hippocampal episodic memory is fundamentally relational, consisting of links between events and the spatial and temporal contexts in which they occurred. Such relations are also important over much shorter time periods, during online visual perception. For example, how do we assess the relative spatial positions of objects, their temporal order, or the relationship between their features? Here, we investigate the role of the hippocampus in such online relational processing by manipulating visual attention to different kinds of relations in a dynamic display. While undergoing high-resolution fMRI, participants viewed two images in rapid succession on each trial and performed one of three relational tasks, judging the images’ relative: spatial positions, temporal onsets, or sizes. As a control, they sometimes also judged whether one image was tilted, irrespective of the other; this served as a baseline item task with no demands on relational processing. All hippocampal regions of interest (CA1, CA2/3/DG, subiculum) showed reliable deactivation when participants attended to relational vs. item information. Attention to temporal relations was associated with more robust deactivation than the other conditions. One possible interpretation of such deactivation is that it reflects hippocampal disengagement. If true, there should be reduced information content and noisier, less reliable patterns of activity in the hippocampus for the temporal vs. other tasks. Instead, analyses of multivariate activity patterns revealed more stable hippocampal representations in the temporal task. Additional analyses showed that this increased pattern similarity was not simply a reflection of the lower univariate activity. Thus, the hippocampus differentiates between relational and item processing even during online visual perception, and its representations of temporal relations in particular are robust and stable. Together, these findings suggest that the relational computations of the hippocampus, known to be important for memory, extend beyond this purpose, enabling the rapid online extraction of relational information in visual perception.

## Introduction

Our perception of the world is not merely a collection of the myriad items in the environment. We do not perceive items in isolation, but rather in terms of their relationship to other items and the context in which they occur. Such relational discriminations are ubiquitous in everyday life. When placing two paintings side-by-side on a wall, we might make fine spatial discriminations to determine whether one is shifted vertically with respect to the other. Knowledge of which of two cars arrived first at an intersection might determine which of them has the right of way. When deciding which piece of fruit to buy at a grocery store, we might find it useful to compare their sizes.

Although relational attention is a key component of visual perception, studies of attention have traditionally focused on perception of individual features or locations (Kastner & Ungerleider, 2000; Maunsell & Treue, 2006). As a consequence, the neural substrates of relational judgments in online visual perception have been largely unexplored (though see Franconeri et al., 2012; Michal et al., 2016). One candidate system for supporting relational attention is the hippocampus, a structure traditionally studied for its role in long-term memory, and particularly for relational forms of long-term memory (Eichenbaum & Cohen, 2014).

One possibility is that the scope of relational processing in the hippocampus is limited to long-term memory, insofar as some have argued that the hippocampus is a dedicated memory system (Squire, Stark, & Clark, 2004). Alternatively, the hippocampus may perform relational computations in a more general way across many domains of cognition (Aly & Turk-Browne, 2018; Olsen, Moses, Riggs, & Ryan, 2012; Yonelinas, 2013), and may therefore also be involved in relational perception. In the current study, we addressed this question by comparing BOLD activity in the hippocampus when attention is directed to individual items vs. to the relations between items. If the hippocampus plays a general role in relational processing, even on the short timescale of perception, its activity should be modulated by the demand to attend to items vs relations.

A second question of interest concerns the types of relational representations the hippocampus might support. Studies implicating the hippocampus in relational memory have largely focused on spatial and temporal processing (Eichenbaum, 2017). For example, the hippocampus is necessary for allocentric spatial navigation (Burgess, Maguire, & O’Keefe, 2002), and contains “place cells” that fire when an animal is in a specific location in the environment (Ekstrom et al., 2003; O’Keefe & Dostrovsky, 1971). The hippocampus also contains “spatial view cells”, which respond to locations that an animal is looking at, even in the absence of navigation (Rolls & Wirth, in press). These findings inspired proposals that the hippocampus is important for memory of spatial context (Burgess, Becker, King, & O’Keefe, 2001; Davachi, 2006; Eichenbaum et al., 2007) and for the construction of spatially coherent scenes (Maguire & Mullally, 2013).

Hippocampal activity also represents the temporal order of experience (Barnett, O’Neil, Watson, & Lee, 2014; Eichenbaum & Cohen, 2014; Kesner & Hunsaker, 2010; Manns, Howard, & Eichenbaum, 2007; Paz et al., 2010; Ranganath, in press; Sakon et al., 2014). For example, during quiet wakefulness and sleep, hippocampal place cells fire in the same sequential order as in previous navigation episodes (Carr, Jadhav, & Frank, 2011). Indeed, some hippocampal cells (“time cells”) fire during successive moments in a temporal delay, keeping a record of elapsed time (MacDonald, Lepage, Eden, & Eichenbaum, 2011; Pastalkova, Itskov, Amarasingham, & Buzsaki, 2008). These studies in rodents converge with work in humans that demonstrates a critical role of the hippocampus in memory for temporal sequences (see Davachi & DuBrow, 2015; Ranganath & Hsieh, 2016).

Although the spatial and temporal representations of the hippocampus have received the most attention, there is also evidence that the hippocampus is critical for other kinds of relational memories as well (Eichenbaum, 2004; Konkel, Warren, Duff, Tranel, & Cohen, 2008; Konkel & Cohen, 2009; McKenzie et al., 2016). For example, Konkel et al. (2008) presented triplets of novel visual objects to patients with hippocampal damage. The object triplets were presented in a particular spatial arrangement and appeared in a particular order. The object triplets were first presented in a study phase, and memory for them was subsequently tested. There were 3 kinds of memory tests, which assessed memory for different kinds of relations between the triplets: spatial, sequential, and associative relations. In the spatial test, patients were shown the objects again and asked to report whether the objects occurred in the same locations during the study phase. In the sequential test, they were to report whether the items were presented in the same sequential order during the study phase. In the associative test, they were to report whether the items were all shown together during the study phase. Patients with damage to the hippocampus performed at chance in all three tasks, suggesting that the hippocampus is critical for relational memory beyond the spatial and temporal domains. Motivated by these studies, we incorporated multiple relational tasks in the current work: in addition to tasks requiring judgments of spatial and temporal relations, we include a task assessing size relations to test the specificity of relational processing in the hippocampus. Importantly, our current study tested *online relational attention*, unlike the Konkel et al., study, which was a test of relational long-term memory.

We collected high-resolution structural and functional MRI data in order to examine the role of different hippocampal subfields in online relational processing. We segmented the hippocampus into subiculum, CA1, and a combined region of interest for CA2, CA3, and dentate gyrus (which cannot be separated at the resolution of our fMRI scans). Below, we describe our predictions for these subfield regions of interest.

First, hippocampal subfield CA1 has been linked to both spatial and temporal processing (Eichenbaum, 2014). For example, CA1 activity codes for the position of items and their spatial context (McKenzie et al., 2014), and tracks changes in the locations of perceived items vs their remembered positions in memory (Duncan et al., 2012). CA1 activity patterns are also modulated by spatial attention (Aly & Turk-Browne, 2016a, 2016b). Moreover, time cells were first discovered in CA1 (MacDonald et al., 2011; Pastalkova et al., 2008); such cells may be important for discriminating the passage of time on the order of seconds or less, as needed in the online temporal attention task in the current study. Indeed, CA1 is necessary for discriminating between memories that were experienced close to one another in time (Gilbert et al., 2001). We therefore predicted that CA1 would be modulated by both spatial and temporal attention.

In contrast to CA1, the literature is mixed on the role of CA3 in processing temporal information, with some (Farovik, Dupont, & Eichenbaum, 2010; Kesner & Hunsaker, 2010; Salz et al., 2016) but not all (Mankin et al., 2012) studies linking this region to the representation of time or sequences. There is evidence that neural activity in CA2 codes for the passage of time, though this might be on the order of hours to days (Mankin et al., 2015). Finally, to our knowledge, no studies have examined the dentate gyrus for potential time cells, but lesions to the dentate gyrus do not impair the ability to make fine temporal discriminations in memory (Gilbert et al., 2001). Because of the mixed evidence on seconds-level timing in CA2, CA3, and dentate gyrus, we made no *a priori* predictions about whether our combined CA2/3/DG region of interest would be modulated by temporal attention. We did, however, expect this region to be modulated by spatial attention (e.g., Aly & Turk-Browne, 2016a, 2016b).

The subiculum plays an important role in spatial processing and navigation (Boccara et al., 2010; Dalton & Maguire, 2017; Hodgetts et al., 2017; Lever et al., 2009; Taube et al., 1990), so we expected it to be modulated by spatial attention (e.g., Aly & Turk-Browne, 2016a, 2016b). To our knowledge, there is very little work on temporal processing signals in subiculum, but there is evidence that subiculum activity patterns are shaped by temporal regularities in experienced events (Schapiro et al., 2012). Thus, it is possible that subiculum will also be modulated by attention to temporal relations.

If the hippocampus plays a role in processing all forms of relations — not just spatial and temporal — we would additionally expect modulation by attention to size relations. We did not have predictions about subfield dissociations for the size task, given the lack of past work exploring this form of relational processing in the hippocampus.

To assess if modulation by online relational attention was specific to the hippocampus, we also examined regions of interest in the surrounding medial temporal lobe (MTL) cortex: parahippocampal cortex (PHC), perirhinal cortex (PRC), and entorhinal cortex (ERC). PHC has been consistently linked to the processing of spatial (Diana, Yonelinas, & Ranganath, 2007; Epstein & Kanwisher, 1998) and temporal (Turk-Browne et al., 2012) context, so we predicted that it would be modulated by attention to both spatial and temporal relations. PRC has been consistently linked to the processing of items (Diana, Yonelinas, & Ranganath, 2007), although recent studies have also linked it to representations of spatial (Bos et al., 2017) and temporal (Naya & Suzuki, 2011) context. Thus, it is possible that PRC will be modulated by attention to both spatial and temporal relations as well. Finally, like CA1, entorhinal cortex (ERC) codes for both spatial and temporal information (Eichenbaum, 2014; Hafting et al., 2005; Kraus et al., 2015; Tsao et al., 2018), so we predicted that this region would be modulated by both spatial and temporal attention.

Although we predict that MTL cortex will be modulated by relational attention, there is an alternative possibility. It has been argued that the hippocampus is unique in forming flexible, relational representations between items (Eichenbaum, Otto, Cohen, 1994). If so, then MTL cortex might only be modulated by attention to items and not modulated by attention to spatial, temporal, or size relations.

To summarize, our approach allows us to test the type and ubiquity of relational representations in the medial temporal lobe: whether such representations exist on the order of seconds during the time-course of perception, how broadly relational computations are applied beyond the spatial and temporal domains, and whether online relational representations are limited to the hippocampus or are also properties of MTL cortex.

## Methods

### Participants

Fifteen participants (8 female, ages 22-33), with normal or corrected-to-normal vision participated for monetary compensation. The study was approved by the Princeton University Institutional Review Board and all participants provided informed consent.

### Stimuli

Stimuli were grayscale images of faces and scenes, equated for luminance. Faces had neutral expressions; half were male and half were female. Half of the scenes were indoor scenes and half were outdoor scenes. Stimuli were presented on a projector screen at the back of the MRI scanner bore and were viewed through a mirror attached to the head coil. We selected stimuli from a pool of 96 faces and 96 scenes, each of which was presented once per run.

Stimuli could be presented within a range of spatial positions on the screen. The center of the reference image on each trial was randomly chosen to be between 0-20 pixels to the left or right of fixation. The reference images also ranged in size on each trial, starting at 81 x 81 pixels and varying up to 10 pixels smaller or larger. Varying the overall position and size of each image increased demands on relational processing (see below), because the relational tasks could not be performed by attending to only one of the images (e.g., by looking for an established larger size if only a single “large” and a single “small” image size had been used). In other words, this variation in absolute position and size was the baseline for relative differences between the two images that served as the basis of the relational tasks, requiring a focus on relative rather than absolute properties.

### Procedure

On each trial, participants were presented with a face and a scene, one above the other (Figure 1). The two items were presented so that one of them was to the left of the other, one appeared first on the screen first, and one of them was smaller than the other. Correspondingly, three relational attention tasks were possible: spatial, timing, and size. In addition, each item could, independently of the other, be tilted clockwise or counterclockwise, which enabled an item attention task that did not require relational processing. After the presentation of each pair, participants were shown a response cue (either black or gray) that pointed up or down, indicating which item should be used as the reference for the task judgment. In the spatial task, participants indicated whether the cued item was to the left of the other item. In the timing task, they indicated whether the cued item appeared first. In the size task, they indicated whether the cued item was smaller than the other. In the item task, they indicated whether the cued item was tilted or not. A post-cueing design was used, instead of a pre-cuing design, because if a cue was presented prior to the onset of a trial, participants would be able to perform the item task by attending to the cued item alone. With a post-cue design, participants had to attend to both items for the relational and the item tasks.

**Figure 1.**
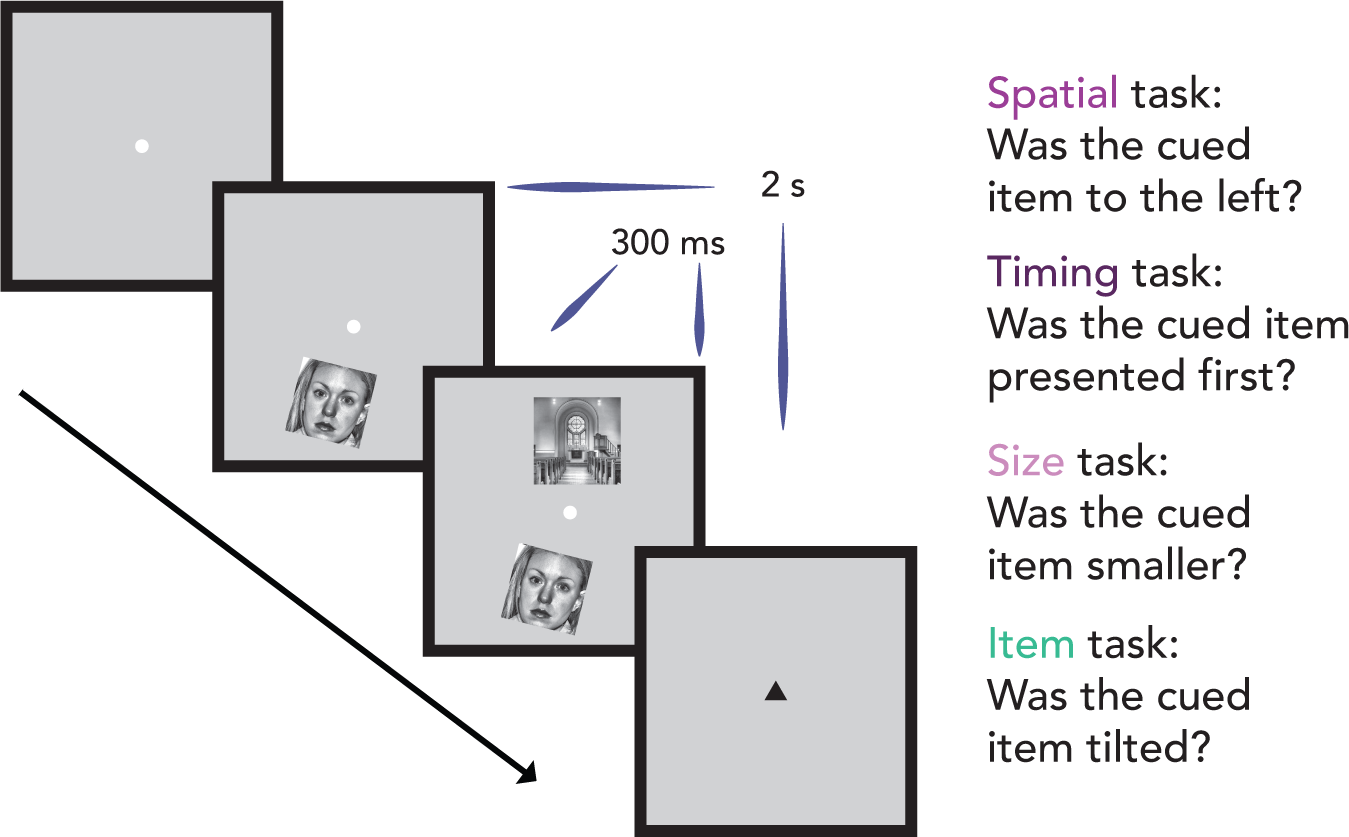
Experimental design. On every trial, participants were presented with a face and a scene, one above and one below fixation. One image was to the left of the other, one appeared on the screen first, and one was smaller than the other. In addition, each item could independently be tilted clockwise or counterclockwise. Participants were cued before every block of 8 trials with the name of the task they were to perform on that block: one of the three possible relational attention tasks (spatial, timing, and size) or the item task. After the presentation of each image pair, participants were shown a response post-cue (either black or gray) that pointed up or down, indicating which item should be used as the reference for the task judgment. Trials lasted 2s in duration, though the two images were shown on the screen for only 300ms.

We included an additional task, in which participants indicated whether the post-cue was black or not, to provide another potential baseline. However, the item task provides a tighter control: as in the relational tasks, the item task required that participants attend to both images presented on each trial (because of the post-cue), with the key difference being that the images could be processed separately and did not need to be judged against each other. Thus, we used the item task as the control against which to examine hippocampal modulation by the relational tasks.

These tasks were completed in a block design, with instructions given prior to each block via on-screen message: “spatial”, “timing”, “size”, “item”, or “cue”.

### Pre-scan behavioral session

To prevent neural differences across tasks from being confounded by differences in task difficulty, we used a staircasing procedure to equate performance across the tasks as much as possible. Each participant completed a behavioral session the day before the scan to staircase performance to 75% accuracy. To reach this performance threshold for each task, we made trial-by-trial adjustments to the relational parameters: spatial separation, temporal delay, and size differences between images for the relational tasks; the degree of tilt of the images for the item task; and shade of gray of the cue for the cue task. A separate staircase was run for each task.

All participants completed one block of 60 trials of each task with initial parameters (Table 1). Participants then completed 4-5 staircased runs. Each run contained one block of 64 trials for each of the 5 tasks, with the order of blocks counterbalanced across runs and with the five image dimensions fully counterbalanced within block (vertical position, category [face or scene], spatial position, size, and tilt of cued item). When participants responded correctly 4 trials in a row, we increased difficulty by one step. If they responded incorrectly on a trial, we decreased difficulty by one step. Participants controlled the onset of trials by pressing a button to continue to the next one.

**Table 1.** Task parameters for the staircasing procedure. The first column indicates the tasks that participants performed. The second indicates the relevant parameter under manipulation. The third contains the initial values of each of the parameters (e.g., images were horizontally offset by 30px with respect to each other). Step size refers to the amount by which we changed the parameter of interest during the staircasing procedure (e.g., whenever participants performed 4 trials of the spatial task correctly in a row, we decreased the horizontal offset by 0.5 pixels). The parameters were not able to go lower than the lower bound (5^th^ column) or above the upper bound (6^th^ column). For the spatial and size tasks, the relevant parameter (position and size, respectively) of the reference image was varied to avoid the possibility that participants could perform the task by attending to only one of the items. The last column shows the variability possible in the reference image. The value for color indicates whether the cue was black (a value of 0) or some shade of gray (a value between 1 and 110).

| Task | Parameter | Initial value | Step size | Lower bound | Upper bound | Reference variability |
| --- | --- | --- | --- | --- | --- | --- |
| Spatial | horizontal offset | 30px | 0.5px | 0px | 150px | -20px – 20px from center |
| Timing | temporal delay | 0.20s | 0.0167s | 0s | 0.2335s | N/A |
| Size | size difference | 30px | 0.5px | 0px | 30px | 151px – 171px |
| Item | degree of tilt | 30° | 1° | 0° | 45° | N/A |
| Cue | color of cue | 100 | 1 | 0 | 110 | N/A |

The final parameters from each participant’s staircasing session (average parameters in **Appendix A**) were then set as the parameters for their fMRI session, with the aim of equating performance across tasks in the scanner as best possible. However, these environments differed, with staircasing conducted on a laptop in a testing room and the fMRI stimuli projected on a screen from a different computer and viewed with a mirror. In particular, the rear-projection display system during fMRI had worse perceived contrast, and so the luminance of the cue had to be adjusted slightly.

### fMRI session

#### Attention tasks

Runs of the attention tasks consisted of an on-off block design, with twelve 16-s blocks of attention tasks (“on”) interleaved with 8-s blocks of fixation (“off”). Task blocks consisted of 8 trials in which a face and scene (identity determined pseudorandomly) appeared above & below fixation. Trial onsets (i.e., the onset of the first item) were time-locked to the repetition time (TR = 2s) and triggered by the scanner. The duration of the stimulus that appeared onscreen first was 300ms. As determined from the staircasing session, there was a spatial offset, temporal delay, and size difference between the images on every trial, and each image could be tilted. Also, as in the behavioral session, we drew from a range of reference spatial positions and image sizes so that the relational tasks could not be performed by only attending to one of the images. Each run contained 12 blocks: 4 blocks of the item task, 4 of the cue task, and 4 of one of the relational tasks. This led to 3 run types — spatial runs, timing runs, and size runs — depending on which relational task was performed. In total, participants completed 384 trials of the cue task and item task, and 128 trials of each of the relational tasks. Participants completed all 4 runs for a particular relational task consecutively. The order of runs was counterbalanced across participants and the order of blocks within a run was counterbalanced within participants.

#### Localizer run

Participants completed a category localizer with alternating blocks of individual faces or scenes. Participants responded with a button box to indicate whether faces were male or female and whether scenes were indoor or outdoor. The structure and timing of the blocks followed the attention task runs (2s trials, 8 trials per block, 16s task/8s fixation, 12 blocks per run). The order of blocks was counterbalanced across subjects. Data from this run were not used in the current study.

### fMRI methods

#### Data acquisition

MRI data were acquired with a 3T Siemens Skyra scanner. Functional images were collected with a gradient-echo EPI sequence (TR = 2000ms; TE = 37ms; FA = 71°; matrix = 128 × 128). Each of 149 volumes contained 27 slices (1.5mm isotropic) perpendicular to the long axis of the hippocampus. The partial-volume images were optimized for hippocampal imaging, and therefore excluded parts of occipital, parietal, and frontal cortices. A high-resolution 3D T1-weighted MPRAGE scan was collected for registration. A high-resolution T2-weighted turbo spin-echo scan (60 slices; 0.4×0.4mm in-plane; 1.5mm thickness) was collected for manual segmentation of hippocampal subfields and MTL cortex.

#### Preprocessing

fMRI data were analyzed with FSL and MATLAB. The first five volumes of each run were discarded for T1 equilibration. All images were skull-stripped to improve registration. The images were preprocessed with motion correction (MCFLIRT), slice-time correction, spatial smoothing (5mm FWHM), high-pass filtering (144s cutoff), and FILM prewhitening.

#### Region of interest segmentation

Manual segmentation of hippocampal subfields and MTL cortex were conducted using published criteria (Aly & Turk-Browne, 2016b; Duvernoy, 2005; Insausti, 1993; Insausti et al., 1998; Mueller & Weiner, 2009; Pruessner et al., 2002; Yushkevich et al., 2010). We segmented these regions of interest (ROIs) on the T2-weighted scans of each participant. MTL ROIs were entorhinal cortex (ERC), perirhinal cortex (PRC), and parahippocampal cortex (PHC). Hippocampal subfield ROIs were subiculum (SUB), CA1, and a combined region for CA2, CA3, and dentate gyrus (CA2/3/DG). All ROIs were traced on coronal slices using FSLview along the entire length of the hippocampus. Sample segmentations, for one anterior slice and one posterior slice, are shown in Figure 2.

**Figure 2.**
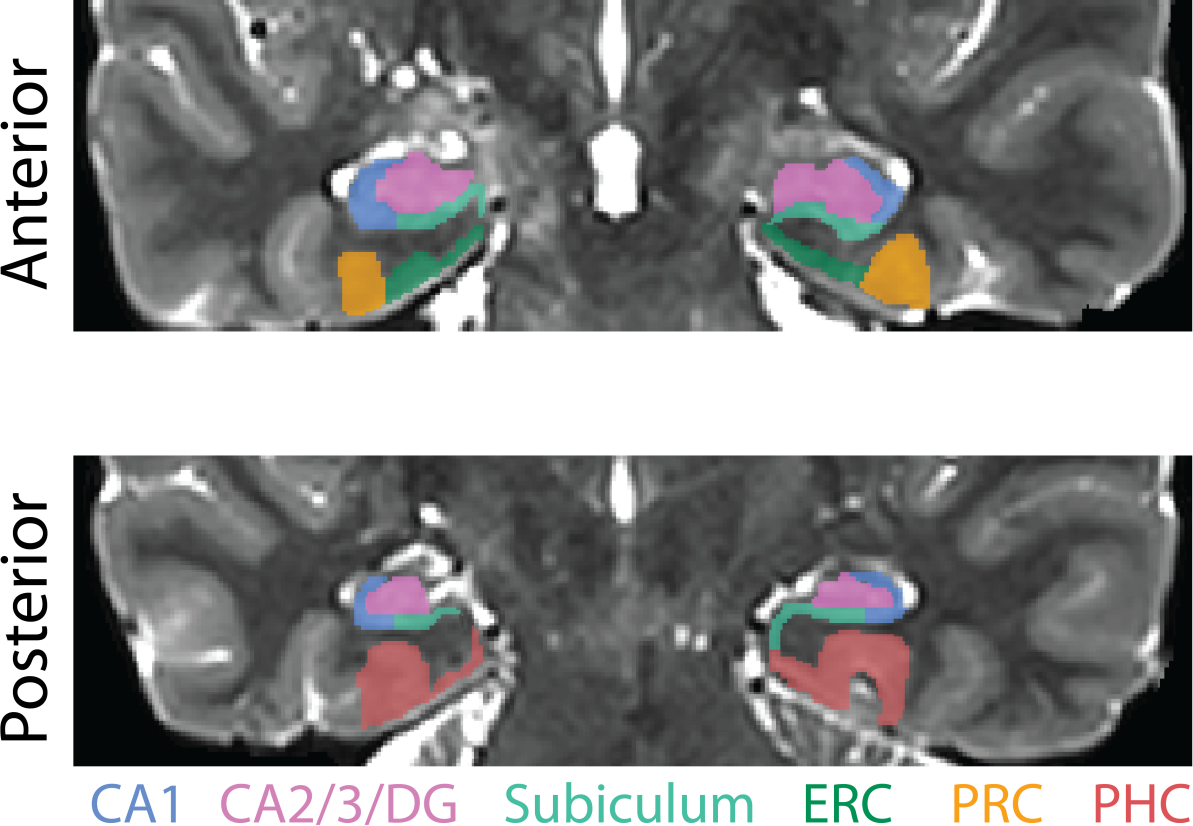
Regions of interest. Example segmentation from one participant is shown for one anterior and one posterior slice. Regions of interest were hand-drawn on individual-participant T2 images. The hippocampal regions of interest were subiculum, CA1, and a combined region of interest for CA2, CA3, and dentate gyrus (CA2/3/DG). The medial temporal lobe cortex regions of interest were entorhinal cortex [ERC], perirhinal cortex [PRC], and parahippocampal cortex [PHC].

The anterior border of PRC was defined as the most anterior slice in which the collateral sulcus (CS) was visible. The posterior border of PRC was the last slice in which the hippocampal head was visible (Poppenk, Evensmoen, Moscovitch, & Nadel, 2013). The lateral border was at the base of the lateral bank of CS. The medial border depended on whether or not ERC was present. For slices without ERC, the border of PRC coincided with the amygdala. For slices with ERC, the medial border was found halfway up the medial bank of CS. PHC was traced from the first slice of the hippocampal body to the last slice of the hippocampal tail. The lateral border of PHC was perpendicular to the lateral bank of the collateral sulcus. The medial border was the border with SUB, perpendicular to the gray matter bend. The anterior border of ERC was found one slice anterior to the start of the frontotemporal junction. The posterior border was the last slice with the hippocampal head. The lateral border was PRC, and the medial border was the border with SUB, perpendicular to the gray matter bend. CA1, CA2/3/DG, and SUB were traced on all slices in which the hippocampal formation was visible. The medial border of the subiculum was ERC in slices containing the hippocampal head and PHC in slices containing the body and tail. At its most anterior slice, the subiculum comprised the entire ventral aspect of the hippocampus (Duvernoy, 2005); the lateral boundary (with CA1) gradually moved medially until, at the body of the hippocampus, the lateral boundary was at the medial edge of the hippocampus at the point where it pinches into a tear shape. CA1 curved around the lateral edge of the hippocampus and bordered CA2/3 at the dorsal aspect of the hippocampus. The boundary between CA1 and CA2/3/DG was determined by the thickness of CA1 on that slice — usually the upper and lateral 2-3 rows of voxels in the hippocampal formation.

#### Univariate analysis

We estimated stimulus-evoked BOLD responses with a general linear model (GLM) containing block regressors convolved with a canonical hemodynamic response function (HRF), which captured the mean evoked response across blocks. Each run was modeled separately in first-level analyses. The 4 runs of the same condition were then combined in second-level analyses. For each condition, we registered the parameter estimate images to the participant’s T2 image, converted the parameter estimates to percent signal change, and extracted the average percent signal change over all voxels in each hippocampal and MTL ROI. We then performed random effects *t*-tests across participants. To isolate signals related to relational processing specifically, we compared evoked activity for each relational task vs. the item task.

#### Multivariate pattern similarity analysis

We combined the first-level analyses for even and odd runs of each type (spatial, timing, size) in a second-level analysis. We then registered the parameter estimate images to the participant’s T2-weighted anatomical image and extracted the parameter estimate for each voxel of every ROI for each of the tasks. To calculate pattern similarity for each task and ROI, we reshaped into vectors the across-voxel activity patterns in each ROI. The pattern similarity score for that task and ROI was the correlation between the vectors from even and odd runs. For example, pattern similarity for the timing task was the Pearson correlation between the mean pattern of activity across voxels for the timing task during odd runs and the mean pattern of activity across voxels for the timing task during even runs. For each participant, task, and ROI, we averaged the pattern similarity measures across the left and right hemispheres of the brain — we did not predict hemispheric differences and therefore this reduced the number of statistical comparisons. Because the item task was included in all three relational run types, we calculated pattern similarity for the item task in each run type separately, and then averaged across run types, resulting in an overall item pattern similarity score. As described above for univariate analyses, to isolate information related to relational processing, we compared pattern similarity for each relational task to the item task.

#### Multivariate-univariate dependence (MUD) analysis

Univariate and multivariate measures are not necessarily independent (Coutanche, 2013; Davis et al. 2014). Indeed, univariate effects (i.e., modulation of overall activity levels) can often drive multivariate ones (i.e., similarity of activity patterns). We therefore quantified the relationship between univariate and pattern similarity measures to assess whether attentional modulation effects in each measure were related or distinct, using an approach known as “multivariate-univariate dependence” (MUD) analysis (Aly & Turk-Browne, 2016a).

The MUD analysis consists of computing the contribution of each voxel to pattern similarity and then calculating the Pearson correlation between these contribution scores and voxels’ level of activity. This quantifies and describes the relationship between univariate activity and pattern similarity: a positive correlation indicates that voxels with the highest activity contribute most to pattern similarity, a negative correlation indicates that voxels with the lowest activity contribute most to pattern similarity, and a zero correlation indicates that a balance of activation and deactivation leads to a stable pattern.

To implement the MUD analysis, we used the same vectors of parameter estimates that we extracted for the pattern similarity analyses. For each participant, ROI, and task, we first normalized the parameter estimates by subtracting the mean and dividing by the root sum-of-squares. We then computed, for each voxel in an ROI, the product of these normalized values from even and odd runs. These products provide a voxel-specific measure of multivariate “influence” — the extent to which a voxel contributed to the pattern similarity measure for that task. Voxels with a positive product (i.e., two positive values or two negative values) contribute to positive pattern similarity, whereas voxels with a negative product (i.e., one positive and one negative value) contribute to negative pattern similarity. Moreover, the magnitude of the product is proportional to the contribution — the larger the product in absolute terms, the greater the “influence”. The *sum* of these normalized products across voxels is equivalent to the Pearson correlation, hence the relationship between the sign/magnitude of the product and the contribution to pattern similarity.

For each voxel, we also obtained the mean level of univariate activity for each task. Finally, we correlated the multivariate influence scores (i.e., the normalized products) with univariate activity across voxels, for each ROI and task. A reliable correlation across participants (whether positive or negative) would suggest that univariate activity and pattern similarity at least partly capture similar information in the data (see Aly & Turk-Browne, 2016a, for simulations that demonstrate the efficacy of this approach).

## Results

### Behavior

We examined reaction times (RTs) and accuracy for each task of interest: the item task, and the spatial, timing, and size relational tasks (Table 2). Inverse efficiency (RT/accuracy; Townsend & Ashby, 1978), a measure of behavior that accounts for speed/accuracy tradeoffs, was matched across relational tasks (*F*(2,28) = 0.51, *p* = 0.60). Because behavioral performance was not different between the relational tasks, a task-difficulty explanation of hippocampal and MTL activity differences between these tasks (see below) is unlikely. That said, the lack of a behavioral difference between the relational tasks is a null effect that we cannot overly interpret, as failure to demonstrate a difference is not strong evidence for equality.

**Table 2.** Behavioral performance. Means are shown with standard deviation in parentheses.

| Task | Accuracy (%) | RT (seconds) | Inverse Efficiency |
| --- | --- | --- | --- |
| Spatial | 74.86 (7.33) | .95 (.16) | .0128 |
| Timing | 78.18 (12.70) | .98 (.16) | .0128 |
| Size | 71.67 (11.15) | .94 (.16) | .0133 |
| Item | 86.00 (6.40) | .89 (.16) | .0104 |

Despite our efforts to balance performance across all tasks, the item task resulted in better performance than the relational tasks (all pairwise comparisons of inverse efficiency, *p* < 0.01). Nevertheless, the item task served as a common baseline across relational tasks and so this cannot explain neural differences between relational tasks.

### Evoked univariate activity

Compared to the item task, the timing task led to deactivation in all hippocampal and MTL cortical ROIs (Figure 3A-F; PHC: *t*(14) = −6.23, *p* = 0.000022; PRC: *t*(14) = −5.61, *p* = 0.000064; ERC: *t*(14) = −4.88, *p* = 0.00024; SUB: *t*(14) = −5.28, *p* = 0.00011; CA1: *t*(14) = −6.20, *p* = 0.000023; CA2/3/DG: *t*(14) = −5.76, *p* = 0.000049). The spatial task was associated with deactivation relative to the item task in the MTL cortical ROIs (PHC: *t*(14) = −3.17, *p* = 0.0068; PRC: *t*(14) = −3.13, *p* = 0.0073; ERC: *t*(14) = −2.54, *p* = 0.024) but not the hippocampal ROIs (SUB: *t*(14) = −1.07, *p* = 0.30; CA1: *t*(14) = −1.76, *p* = 0.10; CA2/3/DG: *t*(14) = −1.84, *p* = 0.09). There were no differences in univariate activity between the size task and the item task in any ROI (PHC: *t*(14) = 0.50, *p* = 0.63; PRC: *t*(14) = −0.20, *p* = 0.85; ERC: *t*(14) = 0.42, *p* = 0.68; SUB: *t*(14) = 0.95, *p* = 0.36; CA1: *t*(14) = 0.47, *p* = 0.65; CA2/3/DG: *t*(14) = 1.26, *p* = 0.23).

**Figure 3.**
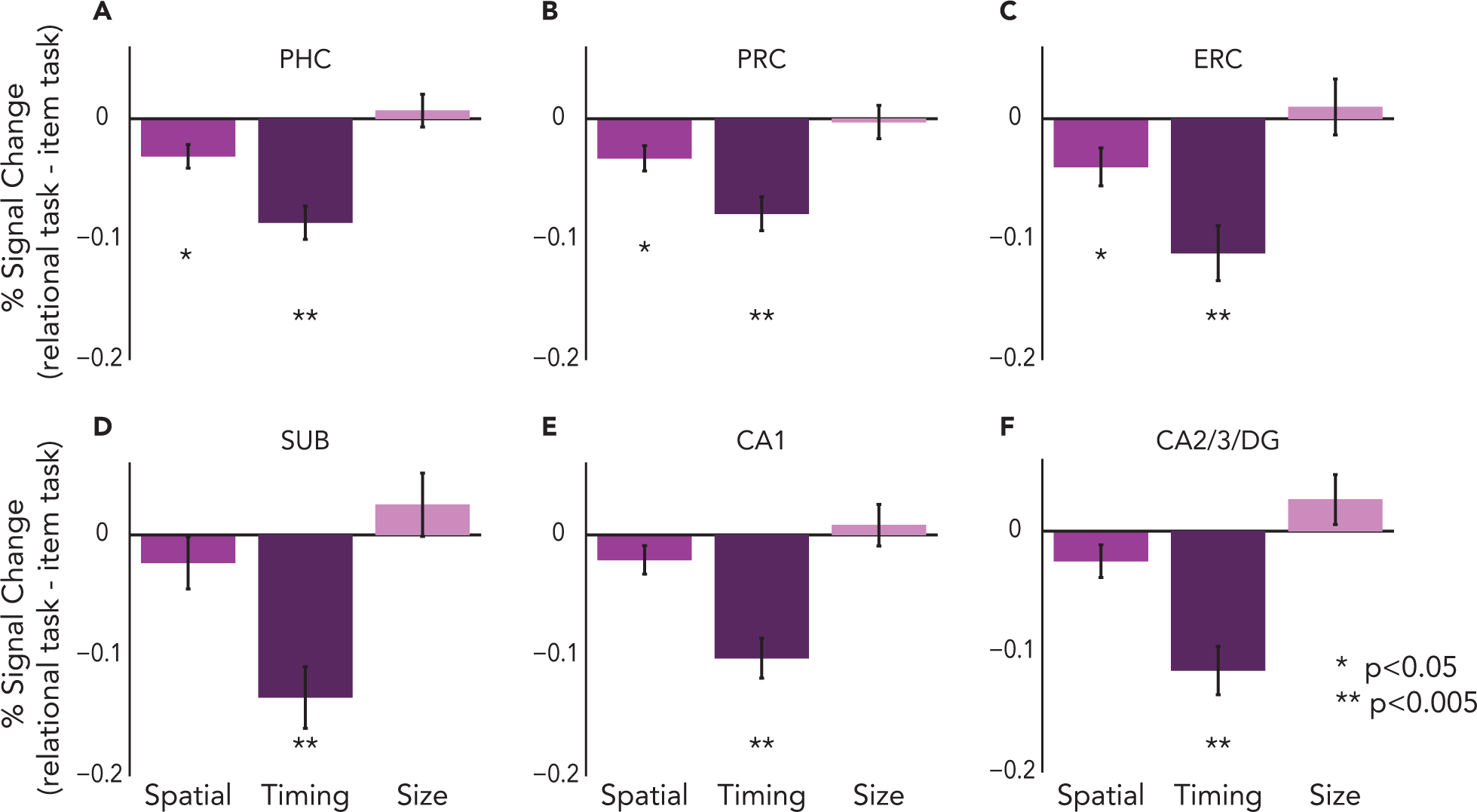
Univariate evoked activity. Percent signal change for the spatial, timing, and size relational tasks, relative to activity for the item task, in each MTL cortical ROI (A: parahippocampal cortex [PHC]; B: perirhinal cortex [PRC]; C: entorhinal cortex [ERC]) and each hippocampal ROI (D: subiculum [SUB]; E: CA1; F: CA2/3/DG). Error bars reflect +/–1 SEM across subjects.

Comparing the relational tasks directly, each of the hippocampal and MTL cortical ROIs showed significant differences (i.e., main effect of relational task, PHC: *F*(2,28) = 16.68, *p* = 0.000017; PRC: *F*(2,28) = 8.73, *p* = 0.0011; ERC: *F*(2,28) = 8.308, *p* = 0.0015; SUB: *F*(2,28) = 12.58, *p* = 0.00013; CA1: *F*(2,28) = 14.28, *p* = 0.000053; CA2/3/DG: *F*(2,28) = 16.18, *p* = 0.000021). Follow-up *t*-tests showed that the timing task was associated with stronger deactivation compared to both the spatial and size tasks in all ROIs (all *p*s < 0.005).

It is important to note that this pattern of results cannot be attributed to our finding that the item task was easier than the relational tasks. For example, one alternative explanation for the timing task deactivation relative to the item task is that more difficult tasks lead to greater suppression of “default mode” processing in the MTL (Greicius, Supekar, Menon, & Dougherty, 2009). However, if we were simply observing default-mode suppression for more difficult tasks, we should have observed more deactivation for *all* relational tasks vs. the item task, because the item task was easier than all relational tasks. This is not the pattern that we observed, particularly for the size task, which often showed numerically *higher* activity levels compared to the item task. Furthermore, after correcting for multiple comparisons, there were no relationships between univariate activity and behavioral performance across participants, further arguing against the notion that deactivation reflects difficulty with the task (although this is a null effect that should not be over-interpreted). Below, we explore another form of the disengagement hypothesis in more detail.

### Multivariate pattern similarity

Does hippocampal deactivation for the timing task reflect disengagement? As noted above, this cannot be based on task difficulty *per se*, given the lack of reliable deactivation for the other relational tasks in the hippocampus. However, perhaps the timing task engages other brain regions, which in turn reduce the need for active hippocampal processing. If the hippocampus is disengaged during the timing task, then representations in the hippocampus should not contain task-related information and should instead be governed by noise or other idiosyncratic, task-irrelevant processing. Accordingly, activity patterns in the hippocampus should be unreliable across repetitions of the timing task and show reduced pattern similarity. Alternatively, reduced mean activity could reflect a sharper, sparser representation of attended information resulting from demands on relational processing supported by the hippocampus, which would in turn be associated with stable patterns of activity (Aly & Turk-Browne, 2016a, 2016b; Kok, Jehee, & de Lange, 2012).

To assess these alternatives, we performed pattern similarity analyses, with a special interest in the timing task because it was the only task associated with robust deactivation in the hippocampus (Figure 4). These analyses revealed more stable patterns of activity during the timing task relative to the item task in SUB (*t*(14) = 2.28, *p* = 0.038) and CA1 (*t*(14) = 3.22, *p* = 0.0062), but no other ROIs (PHC: *t*(14) = 0.64, *p* = 0.53; PRC: *t*(14) = 1.40, *p*= 0.18; ERC: *t*(14) = 1.50, *p* = 0.15; CA2/3/DG: *t*(14) = 1.48, *p* = 0.16).

**Figure 4.**
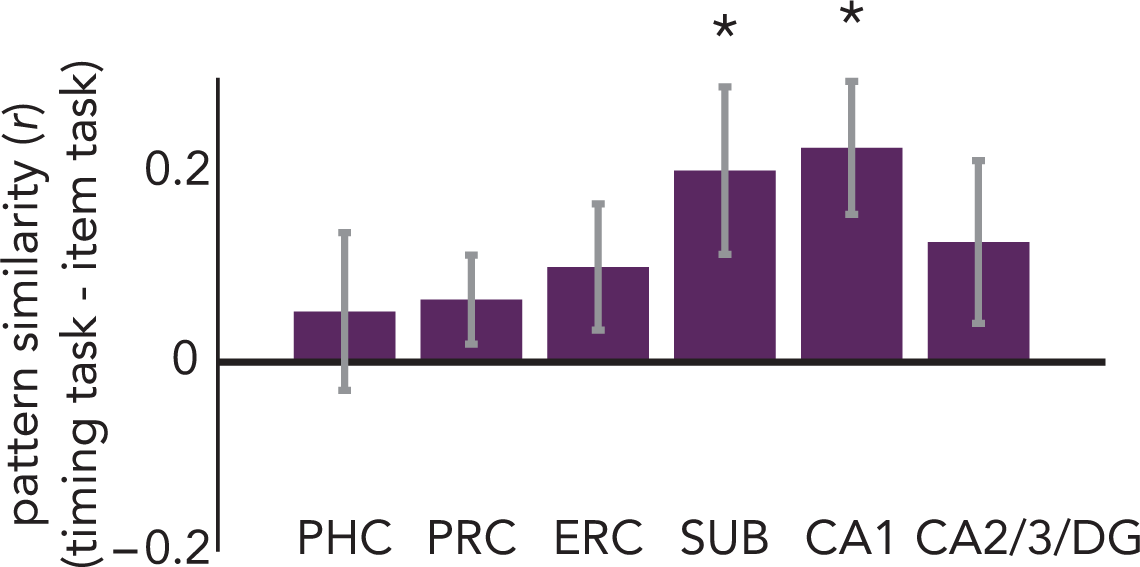
Multivariate pattern similarity for the timing task. Pearson correlation between activity patterns in each ROI for odd vs. even runs of the timing task, relative to the correlation between activity patterns for the item task. Subiculum [SUB] and CA1 showed greater pattern similarity for temporal attention vs. item attention. Error bars reflect +/–1 SEM across subjects. * *p* < 0.05

In contrast, there were no differences between pattern similarity for the spatial task vs item task or the size task vs item task in any hippocampal or MTL cortical ROI (Spatial vs Item: PHC: *t*(14) = 0.20, *p* = 0.84; PRC: *t*(14) = 0.57, *p* = 0.58; ERC: *t*(14) = 0.98, *p* = 0.34; SUB: *t*(14) = −0.10, *p* = 0.92; CA1: *t*(14) = 0.80, *p* = 0.44; CA2/3/DG: *t*(14) = −0.57, *p* = 0.57; Size vs Item: PHC: *t*(14) = 0.47, *p* = 0.65; PRC: *t*(14) = −0.74, *p* = 0.47; ERC: *t*(14) = −1.76, *p* = 0.10; SUB: *t*(14) = 0.17, *p* = 0.86; CA1: *t*(14) = 2.06, *p* = 0.06; CA2/3/DG: *t*(14) = 0.59, *p* = 0.57).

These data suggest that reduced hippocampal activity (in subiculum and CA1) in the timing task may reflect sharpening of representations rather than disengagement from the task. The subiculum effect, however, was weak and did not survive correction for multiple comparisons across regions (FDR corrected *p* = 0.114). In contrast, the CA1 effect is robust even after correcting for multiple comparisons (FDR corrected *p* = 0.037). Because we were interested in the timing task in particular (given the univariate results), we corrected for multiple comparisons across regions of interest, but not across the relational tasks.

### Multivariate-univariate dependence analysis

Subiculum and CA1 showed lower levels of activity for the timing task (Figure 3D-E) but higher pattern similarity for the timing task (Figure 4). Is the increase in pattern similarity a result of univariate deactivation in these ROIs (see Coutanche, 2013)? For example, is the stability of the activity pattern simply a consequence of some voxels consistently deactivating in the timing task, or does the pattern reflect information that is not captured in terms of a mean response? To address this question, we conducted a multivariate-univariate dependence (MUD) analysis to quantify the relationship between univariate activity levels in each voxel and its contribution to pattern similarity (Figure 5; Aly & Turk-Browne, 2016a). Insofar as deactivation is responsible for pattern similarity, we should observe a negative relationship in the MUD analysis: voxels with the lowest activity levels should be the largest contributors to pattern similarity. However, there was no reliable relationship between voxels’ univariate activity and their contribution to pattern similarity for the timing task in SUB (*t*(14) = 0.81, *p* = 0.43) or CA1 (*t*(14) = 0.61, *p* = 0.55).

**Figure 5.**
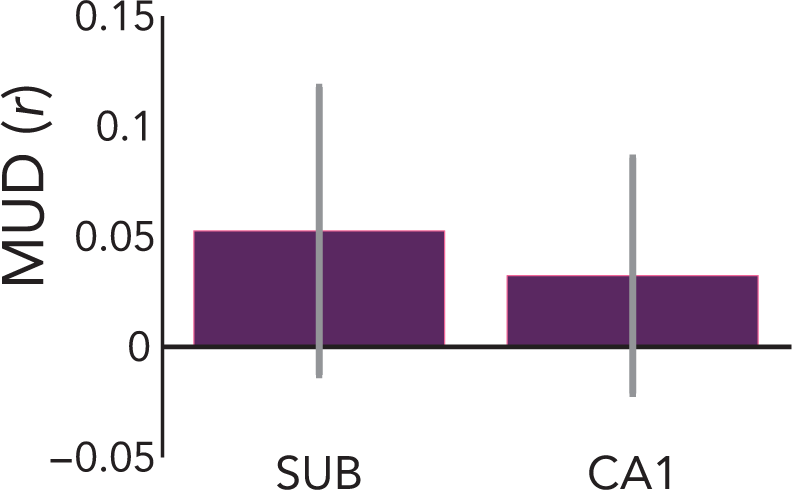
Multivariate-univariate dependence analysis for the timing task. The contribution of each voxel to pattern similarity was estimated by normalizing BOLD activity over voxels within each ROI, separately for the timing task in even and odd runs, and computing pairwise products across runs. To estimate multivariate-univariate dependence, these products were then correlated with the average univariate activity in the timing task for each voxel. Neither SUB nor CA1 showed a relationship between the two measures, suggesting that deactivated voxels were not solely responsible for increased pattern similarity. Error bars reflect +/–1 SEM across subjects.

The absence of a correlation between univariate activity and contributions to pattern similarity across voxels suggests that it is neither high nor low univariate activity that is driving pattern stability: instead, the elevated pattern similarity in the timing task reflects a balance of voxel activation and deactivation that together underlie the stable pattern (see simulations in Aly & Turk-Browne, 2016a, for a demonstration). However, this interpretation rests on a null result (the absence of a relationship between univariate activity and pattern similarity); thus, a Type II error cannot be ruled out. Future studies using the MUD analysis will be important for characterizing when the hippocampus does, and does not, show dependencies between univariate activity and multivariate pattern similarity.

## Discussion

Attention has been studied primarily in terms of individual features and locations, but our experience of the world is fundamentally relational, consisting of representations of items and their associations to other items and the global context. We examined the neural substrates of relational attention, focusing on the hippocampus because of its critical role in relational forms of long-term memory (Cohen et al., 1999; Eichenbaum & Cohen, 2014; Ryan, Althoff, Whitlow, & Cohen, 2000). According to some theories (Squire et al., 2004), the hippocampus is a dedicated memory system, and thus its role in relational processing should be limited to relational memory. Alternatively, the relational computations of the hippocampus might support a more general function, and contribute to relational processing across domains of cognition from perception to long-term memory (Aly & Turk-Browne, 2018; Shohamy & Turk-Browne, 2013; Yonelinas, 2013). In the current study, we tested the hypothesis that the hippocampus would be recruited by relational attention even during online perceptual processing, with no demands on long-term memory. Moreover, we tested whether the hippocampus is specialized for some types of relational processing (e.g., spatial or temporal), or plays a broader role in other types of relations as well (e.g., relative size).

We found strong deactivation throughout the hippocampus when participants attended to temporal relations, as compared to attending to items. This reduction in univariate activity was accompanied by an increase in multivariate pattern similarity in the timing task relative to the item task. These results echo other findings showing that reductions in activity can be accompanied by increases in information content in patterns of activity (Aly & Turk-Browne, 2016a, 2016b; Bell, Summerfield, Morin, Malecek, & Ungerleider, 2016; Kok et al., 2012), and raises the possibility of sparse, sharper representations in the hippocampus when attention is directed to temporal relations. Further analyses indicated that higher pattern similarity in the timing task was not simply a consequence of lower levels of activity: instead, the stable patterns of activity in the timing task were a result of a balance of activation and deactivation. Finally, the selectivity of this pattern of results to temporal attention is unlikely to be due to differences in task difficulty, because all relational tasks were matched in behavioral performance.

We also found that entorhinal, perirhinal, and parahippocampal cortices were all modulated by both spatial and temporal relational attention. However, these regions were *deactivated* for relational vs item attention, and did not show more stable patterns of activity for relational vs item attention. Thus, the current study cannot rule out that these regions were simply disengaged during the relational attention tasks, and this finding requires further investigation.

### What is represented in the hippocampus during online attention?

What is the content of the sharpened representations in the hippocampus during temporal attention? One possibility is that these stable activity patterns represent a specific, yet abstract, attentional state. Another possibility is that these stable activity patterns do not represent an abstract attentional set, but rather represent precise information about the components of stimuli that are attended on the temporal attention trials (e.g., a short-term representation of whether the upper vs lower part of the screen changed first). Our current results cannot adjudicate between these possibilities, because both components (the abstract attentional set, and the precise features that are attended) are inherent aspects of the attention task. Indeed, it is difficult to conceive of any study design that can separate the brain’s representation of an abstract attentional state from its representation of the attended features, because the attended features are a key aspect of defining the attentional state in the first place. What can be concluded, however, is that the patterns of activity in the hippocampus for temporal attention code for the commonalities of that attentional state across different trials that vary in terms of the visual images presented and their precise timing.

Are the results of the current study merely reflecting relational long-term memory in the hippocampus? We think that this is unlikely for several reasons. For example, one might argue that participants are incidentally encoding the stimuli into memory. While this is certainly possible, that is not enough to explain our results. If memory were the driving force behind the differential hippocampal modulation we observed across tasks, then there must have been different amounts of incidental encoding in these tasks — but there is no reason why that should be true. Even if different amounts of incidental encoding were occurring across tasks, our dissociation between univariate activity and pattern similarity in the hippocampus complicates the interpretation. Specifically, greater pattern similarity (in the timing task) and greater univariate activity (in the other tasks) have both been linked to better memory encoding in the hippocampus (Carr et al., 2013; Wolosin, Zeithamova, & Preston, 2013).

Thus, our results cannot be accounted for by appealing to long-term memory, and instead concur with recent neuropsychological and neuroimaging studies highlighting a role for the hippocampus in online processing without demands on long-term memory, including visual perception and attention tasks (e.g., Aly et al., 2013; Aly & Turk-Browne, 2016a; Lee et al., 2012; Warren, Duff, Tranel, & Cohen, 2011; Zeidman & Maguire, 2016; Zeidman, Mullally, & Maguire, 2015). These findings challenge the traditional perspective of the hippocampus as a dedicated declarative memory system (Squire et al., 2004; Squire & Wixted, 2011), and highlight the reach of the hippocampus to attention and perception.

### Space and time in the hippocampus

Along with long-term memory, studies of the hippocampus have also focused extensively on its role in representing space (Bird & Burgess, 2008; Bussey & Saksida, 2005; Kumaran & Maguire, 2005; Nadel, 1991; O’Keefe & Nadel, 1978). Recently, however, there has been increased focus on the importance of the hippocampus for temporal processing — both its contribution to the temporal organization of memories (Davachi & DuBrow, 2015; Eichenbaum, 2013; Howard & Eichenbaum, 2013; Hsieh, Gruber, Jenkins, & Ranganath, 2014; Jenkins & Ranganath, 2010; Paz et al., 2010; Schapiro et al., 2016; Staresina & Davachi, 2009), as well as the perception of time (Barnett, O’Neil, Watson, & Lee, 2014; Palombo, Keane, & Verfaellie, 2016). For example, the hippocampus contains “time cells” whose successive activity signals the passage of time (Eichenbaum, 2014; MacDonald et al., 2011; Pastalkova et al., 2008). Beyond space and time, some studies support the view that the hippocampus engages in relational processing irrespective of content (Eichenbaum, 2004; Hannula, Tranel, & Cohen, 2006; Konkel et al., 2008; McKenzie et al., 2016; Schiller et al., 2015). Most previous work with fMRI, however, has focused on one aspect of relational processing (e.g., spatial relations) and thus has not been in a position to test these different perspectives. By comparing multiple relations in the same experiment (see Konkel et al., 2008 for a similar approach in a patient study of memory), our findings were able to highlight a special role for the hippocampus in the online processing of temporal relations.

We observed reliable deactivation throughout all hippocampal subfields during attention to temporal relations. However, pattern similarity was higher during temporal attention in subiculum and CA1, but not CA2/3/DG. One might have expected stable activity patterns for temporal attention in CA2/3/DG, given that CA3, like CA1, contains time cells (Salz et al., 2016), and is implicated in temporal and sequential processing more generally, including on the order of seconds (Farovik et al., 2010; Kesner & Hunsaker, 2010). However, our region of interest also includes CA2 and dentate gyrus, which have not yet (to our knowledge) been implicated in seconds-level timing (see Mankin et al., 2015 for timing on the order of hours to days; also see Gilbert et al., 2001). Thus, the lack of pattern stability for temporal attention in CA2/3/DG is inconclusive and requires further investigation with studies that can separately examine CA3, CA2, and DG.

Likewise, the lack of an effect for spatial relational attention in the hippocampus is unexpected given extensive evidence that the hippocampus is involved in spatial processing (Eichenbaum & Cohen, 2014), including spatial relational attention (Aly & Turk-Browne, 2016a, 2016b). One possibility is that the hippocampus is involved in fine relational discriminations (e.g., Aly et al., 2013; Barnett et al., 2014) but is not required when such discriminations can be solved on the basis of individual featural comparisons (Baxter, 2009; Bussey & Saksida, 2005). However, we designed the tasks so that such feature-level comparisons would not be sufficient to support performance — that is, the jitter in spatial location and size was meant to ensure that attention to, and comparison of, both items was required to do the relational tasks. Thus, another possibility is that the hippocampus only becomes involved in spatial processing given sufficiently complex or “high-resolution” task demands (Aly et al., 2013; Yonelinas, 2013). For example, the hippocampus is engaged by the demand to attend to, and find similarities or differences in, spatial layouts of complex, naturalistic scenes (Aly & Turk-Browne, 2016a, 2016b; Aly et al., 2013; Lee et al., 2012). It is possible that our spatial relational task does not sufficiently tax the abstract and flexible spatial representations of the hippocampus, and can be solved on the basis of relational representations elsewhere.

This raises an important question for future research: Does the hippocampus become involved at the same level of complexity when assessing relations in the spatial and temporal domains? Or do some types of relations require the hippocampus at an earlier level of complexity than others? These are difficult problems to solve, but these limitations point to a need in the field: a need to define what is “complex” or “high resolution” with enough precision that these definitions can be used to generate testable hypotheses about when the hippocampus should, and should not, be involved in any given task.

### Transformation of relational representations from perception to memory

In studies of long-term memory encoding and retrieval, the hippocampus shows greater univariate activity for relational memory vs item memory (see Davachi, 2006 for review). This contrasts with our current findings, in which the hippocampus showed less univariate activity for relational vs item attention. An open question is why the direction of the relational vs item effect switches from perception to memory. There are at least three possibilities.

The first possibility is that there are two different relational computations in the hippocampus. One supports in-the-moment attention and perception, is sharply tuned, and is associated with reductions in univariate activity. The other supports long-term memory, is more integrative, and is associated with enhancements in univariate activity.

A second possibility is that the same set of relational computations in the hippocampus is expressed in different ways in perception and memory: the initial representation is sharp and sparse, but over time is transformed to a more integrated representation. We believe that this second possibility is unlikely, because the hippocampus shows greater univariate activity for relational vs item *encoding*, arguing against the emergence of a univariate activity enhancement over the time course of consolidation.

A final possibility lies somewhere in between: the hippocampus has a common set of relational computations, but these computations are differently taxed during perception and memory. In attention and perception tasks, stimuli are often repeated many times, presented for a short duration, and only have to be processed enough to accomplish the task in the immediate moment. In memory tasks, a stimulus might be shown only once, and elaborative processing is helpful during both encoding (to create a distinct memory representation) and retrieval (to bring to mind associated details). These task demands might prioritize a finely-tuned, sharp, and short-lasting representation during attention/perception tasks, and a richer, more integrated, longer-lasting representation during memory tasks.

Importantly, this viewpoint suggests that it is not perception vs memory *per se* that produces these differences in the properties of hippocampal representations, but the demands of the typical tasks used to study these cognitive processes (see Aly & Yonelinas, 2012, for a similar perspective). This perspective predicts that making a perception (or attention) task more like a memory task would yield greater hippocampal activity for relational vs item perception (or attention). This might be accomplished by more closely matching perception tasks to the encoding phase of long-term memory studies; for example, by showing stimuli only once each, and for a longer duration. Including a long-term memory test following the perception task would enable separate examination of hippocampal effects related to memory encoding vs those related to perceptual processing (e.g., as in Lee, Brodersen, & Rudebeck, 2013). These and other approaches will be useful for characterizing relational representations in the hippocampus during perception and memory, and determining whether a single set of underlying computations supports both.

## Conclusions

Attention gates what we perceive and remember, and yet we know relatively little about how attention modulates neural activity in the hippocampus. Recent work has made important progress in elucidating how the hippocampus is modulated by the focus of attention (Aly & Turk-Browne, 2016a, 2016b, 2017), in line with the current findings. We provide evidence that the hippocampus is differentially involved in relational and item attention, even during online visual perception. Attention to temporal relations reduces hippocampal activity and increases hippocampal pattern stability, with balanced activation and deactivation producing a sharpened representation. These findings show that the relational computations of the hippocampus can extend beyond long-term memory, enabling the rapid online extraction of relational information during visual perception.

## Appendix A

**Appendix A.**
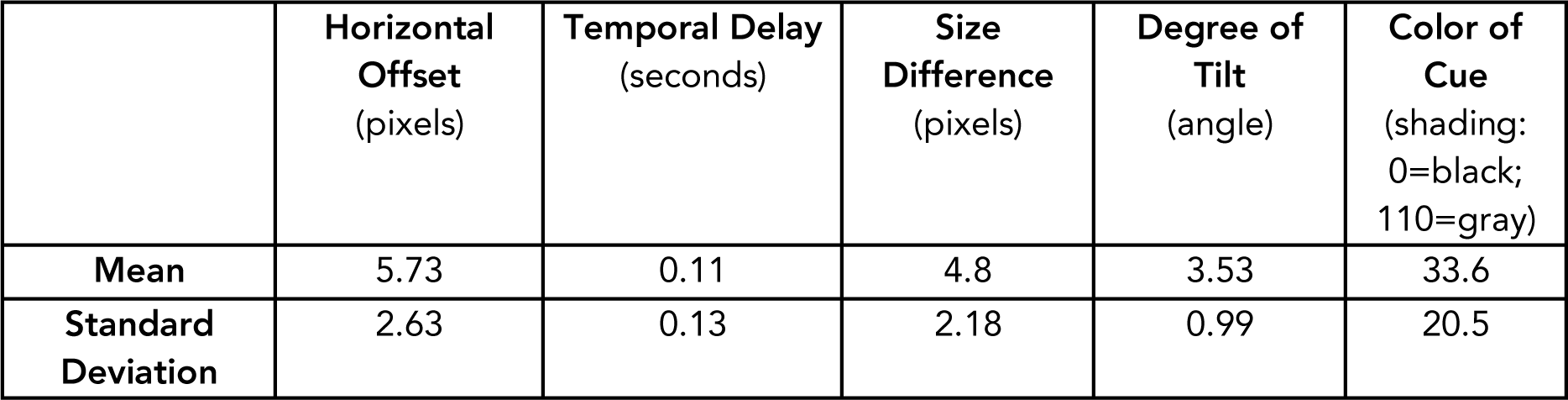
Mean and standard deviation of final parameters from each participant’s staircasing session.

|  | **Horizontal**<br>**Offset**<br>(pixels) | **Temporal Delay**<br>(seconds) | **Size**<br>**Difference**<br>(pixels) | **Degree of**<br>**Tilt**<br>(angle) | **Color of**<br>**Cue**<br>(shading:<br>0=black;<br>110=gray) |
| --- | --- | --- | --- | --- | --- |
| **Mean** | 5.73 | 0.11 | 4.8 | 3.53 | 33.6 |
| **Standard**<br>**Deviation** | 2.63 | 0.13 | 2.18 | 0.99 | 20.5 |

